# Biologically-Constrained Spiking Neural Network for Neuromodulation in Locomotor Recovery after Spinal Cord Injury

**DOI:** 10.1101/2024.09.04.611123

**Authors:** Raymond Chia, Chin-Teng Lin

## Abstract

Presynaptic inhibition after spinal cord injury (SCI) has been hypothesised to disproportionately affect flexion reflex loops in locomotor spinal circuitry. Reducing gamma-aminobutyric acid (GABA) inhibitory activity increases the excitation of flexion circuits, restoring muscle activation, and stepping ability. Conversely, nociceptive sensitisation and muscular spasticity can emerge from insufficient GABAergic inhibition. To investigate the effects of neuromodulation and proprioceptive sensory afferents in the spinal cord, a biologically constrained spiking neural network (SNN) was developed. The network describes the flexor motoneuron (MN) reflex loop with inputs from ipsilateral Ia- and II-fibres and tonically firing interneurons. The model was tuned to a baseline level of locomotive activity before simulating an inhibitory-dominant and body-weight supported (BWS) SCI state. Electrical stimulation (ES) and serotonergic agonists were simulated by the excitation of dorsal fibres and reduced conductance in excitatory neurons. ES was applied across all afferent fibres without phase- or muscle-specific protocols. The present study describes, for the first time, the release of GABAergic inhibition on flexor MNs as a potential mechanism underlying BWS treadmill training. The results demonstrate the synaptic mechanisms by which neuromodulatory therapy tunes the excitation and inhibition of ankle flexor MNs during locomotion for smoother and more coordinated movement.

**Author Summary:** SCI is a life-altering condition that often leaves young adults paralysed and reliant on others for support. Restoring the ability to walk is a critical goal to help improve independence and quality of life for people living with SCI. Promising new treatments, such as spinal cord stimulation and drug therapies, aim to reawaken the neurons that control walking. However, scientists still do not entirely understand how these treatments work. In this study, we developed a detailed computer model of the neural circuits involved in walking to test how therapies such as serotonin-boosting drugs, ES, and BWS training might help. Our findings suggest that these treatments can work together to reduce excessive inhibition that blocks ankle movement, leading to smoother and more coordinated steps. This research helps uncover how these therapies work and provides insights to develop better rehabilitation strategies for improving walking after SCI.

## Introduction

SCI globally affects an estimated 9 million people as of 2019, with an age standardised incident rate of about 109 per 100,000 (1). In the event of SCI, damage to nervous tissue can result in loss of voluntary control, sensation, spasticity, diaphragm dysfunction, pressure ulcers, and pain syndromes (2, 3). Sufferers of SCI often report non-physical symptoms such as emotional disorders, loss of independence, depression, anxiety, and clinical levels of stress (4). The lifelong management places intense financial burden not only the patients and their communities but also on the broader economic landscape (5, 6). Lifetime medical costs in Canada can range from $1.47 to $3 million CAD (2013 prices) per person (7), £1.12 million per person in the UK (2016 prices) (8), and range from $0.77 to $1.3 million USD (1995 prices) in the US (9). Recovering voluntary muscle activity and returning locomotion activity to SCI sufferers could save societal and patient costs while also improving the patient’s quality of life (10, 11).

Recovering gait remains as a top priority for people living with SCI (12). Flexor activity is critical for step progression during locomotion, acting as shock absorbers before foot strike (13), adapting step-height to continue locomotion progression (14), and resetting locomotion (15). Increasing excitability of locomotor networks after paralysis can improve locomotor capabilities however, hyperexcitation of flexor muscles can result in spastic muscle expression, leading to poor balance and coordination (16–18). Maintaining an excitation-inhibition balance of excitability emerges as an intuitive solution to enabling robust locomotor expression.

SCI interrupts normal bidirectional signalling, leading to dysfunctional neural circuitry, tilting the balance of excitation and inhibition (19). Lack of descending activity keep MNs at a predominantly inhibited state while inhibitory populations in the dorsal and intermediate zone become over-reactive (20, 21). A large percentage of the SCI population experience spastic muscle activity, likely due to insufficient release of GABA neurotransmitters (22–26). Nevertheless, even with an overly excited or inhibited environment and detached from brain inputs, the locomotor spinal circuit can continue to express coordinated motor function given sufficient excitation and contextually relevant sensory information (27–31).

Proprioception is a critical sensory input to entrain and recover locomotion after SCI (32–34). Proprioceptive afferent innervation is widespread and diverse, projecting to MNs (35–37), GABAergic (38, 39), and serotonergic (40–43) interneurons (INs) in the dorsal and intermediate zone of the spinal cord. Long and short axons spread across multiple segments and organised spatially and by modality (37, 44– 46). Proprioceptive interneurons (PINs) are mainly excitatory, with most inhibitory populations projecting ipsilaterally (47). Due to their complex and integratory nature, PINs have been suggested to be a possible neural detour around spinal lesions, recovering voluntary sensorimotor control after SCI (46, 48–51).

After SCI, axons spared from injury allow voluntary activation and sensation of the body past lesion sites (51, 52). Traditional rehabilitation therapy leverages these residual connections to maximise motor skills via therapeutic exercise or overcome losses with assistive devises (28, 53). Neuromodulation techniques such as spinal cord ES (30, 31, 54, 55) and pharmacology (51, 56, 57) have shown to recover locomotor activity after SCI. Moreover, chronic application of ES in conjunction with physical rehabilitation enabled volitional muscle activation even without ES (29). Although these observations show promise for new and effective neurorehabilitation therapies, the mechanisms of action and synergy between sensory ensembles and ES remain in question (58, 59). It is natural to seek methods to return excitation to sublesional networks after losing descending input (60). Most ES techniques have sought to excite and entrain locomotion by activating dorsal roots in the epidural space (59, 61–64). However, ES, in the same anatomical space, using similar stimulation protocols, can equally evoke inhibition and restore an overly excited network (23, 24, 65). A more in-depth and nuanced view of ES therapy is required to fully appreciate the complexity of modulating the neural environments. A key question remains: How do neuromodulation therapies integrate with sensory information? We hypothesise that spinal cord locomotor circuits require balanced excitation and inhibition to coordinate flexor activity.

This study describes a biologically constrained tibialis anterior (TA) SNN rodent model receiving heterogeneous excitatory and inhibitory synapses, including GABAergic presynaptic inhibition. A combination of ES and serotonin agonist (5-HT) neuromodulators are simulated in an SCI and SCI with BWS locomotion setting. We show that combining BWS with ES reduces overactive stance-evoked GABA inhibition and returns TA MN firing rates towards baseline.

## Methods

A biologically constrained SNN was developed to investigate the effect of neuromodulation on sensory-driven rodent spinal locomotor circuits. Simulations were run on an Intel Xeon Gold 6238R 2.2GHz Processor. The software was developed in Python 3.10.0 using the Brian2 neural simulator module (v2.6.0) (66). The simulation time step was set to 5 µs and Euler approximations were used for ordinary differential equation solving. A total of 9 steps were simulated, where gait stance and swing phases were split at 65% of the gait cycle (67). This study simulated three different neurological environments, including a baseline, SCI, and SCI with a BWS state. Each neurological state was modulated with inputs from ES and 5-HT.

The SNN model architecture was biologically constrained using synaptic connections inferred from previous cell staining (39, 61, 68, 69), electrophysiological (70–76), and genetic works (77–82). A second biological constraint was set by matching neuron cell dynamics to experimentally measured electrophysiological neural recordings (62, 68, 75, 77, 78, 80, 83–85). Specific parameter settings are described in the below sections and set to be within biologically plausible ranges. Finally, propriosensory inputs were constrained to previously validated musculoskeletal and muscle spindle models (62).

The simulated data was considered dependent and tested using the Shapiro-Wilk method, where *p >* 0.05 was considered a normal distribution. Normal distributions were tested for significance using Tukey Honestly Significant Difference (HSD), and non-normal distributions were tested using the Wilcoxon signed-rank test. The distributions were considered significantly different if *p <* 0.05. Population firing rates were averaged with a 25 ms Gaussian window. Experiment code can be found at https://github.com/rchia16/balancing-locomotor-networks.git.

### Afferent Signal Inputs

Ia and II TA and gastrocnemius medialis (GM) muscle afferent signals were calculated by using musculoskeletal and muscle spindle models during locomotion, described in previously validated computational models (62). To emulate BWS afferent signals, both TA and GM Ia and II data were offset by a scalar amount using values from electrophysiological experiments (86). The new BWS equations were set to equations (1) and (2) where *K*_*GM*_ = *−* 0.6 and *K*_*T A*_ = *−* 0.122, scaling the EMG envelope to 40% and 87.8% respectively, simulating 60% effective body weight. The equations refer to *x* as stretch, *v* as stretch velocity, and *EMG*_*env*_ as the min-max normalised EMG envelope (87). Since *EMG*_*env*_ magnitude ranged between 0 and 1, a scalar offset can be applied to the afferent signal inputs (62). Afferent signals were set as timed array poisson distributed inputs with a sampling frequency of 200 Hz and connected to leaky integrate and fire (LIF) axon models, see equation (3) and Fig. 1 A and B. Parameters were set according to previously validated computational models (61) and tuned to replicate the input firing rate, see table 1. Background noise of afferent axons (*I*_*noise*_) was modelled as a normally distributed variable with a standard deviation scaled to 0.3 pA. Tuning was validated with Pearson correlation coefficient and mean absolute error. For detailed results, refer to table 5 in the Supplementary Information.

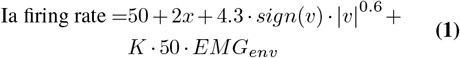

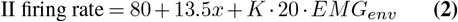

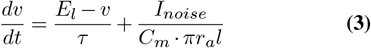

**Table 1.** Afferent axon parameters for LIF model.

| Parameter | Value |
| --- | --- |
| N | 60 |
| $\tau$ | 30 ms |
| $\tau_{refractory}$ | 1.6 ms |
| $E_L$ | -80 mV |
| $V_{th}$ | -60 mV |
| $V_{rest}$ | -70 mV |
| $C_m$ | 0.1 μF/cm<sup>2</sup> |
| $r_{Ia}$ | 9 ± 0.2 μm |
| $r_{II}$ | 4.4 ± 0.5 μm |
| $l$ | 1 mm |

**Fig. 1.**
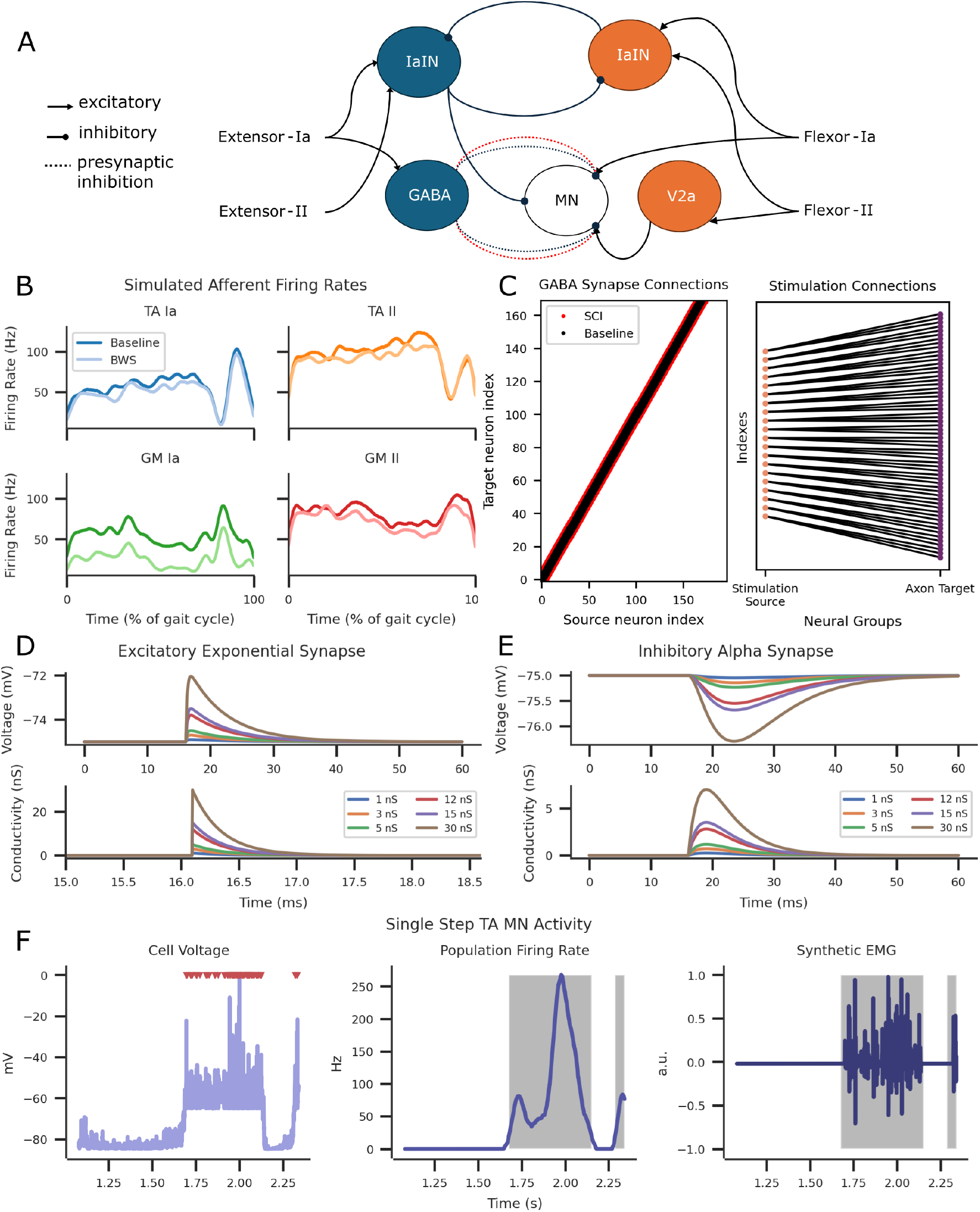
Computational SNN model of the flexor network with GM extensor and TA flexor proprioceptive Ia and II inputs. **A** Biologically constrained SNN ankle flexor model. Arrow ends indicate excitation, circle ends indicate inhibition, and dotted line with circle ends indicate presynaptic inhibition connections. The red dotted lines represent increased SCI induced GABA synapses. **B** Afferent axon firing rates for extensor (GM) and flexor (TA) Iaand II-fibres in the baseline and BWS condition. **C** GABA synapse connections (left) between the source and target neuron in the Baseline and SCI condition. Poisson stimulation input to afferent axon connections (right) during ES condition. SCI synapses retained the Baseline connections with the addition of the synapses depicted in red. **D** Excitatory exponential synapses and **E** inhibitory alpha synapses across different conductances. **F** Simulated TA MN output from a single step. Left illustrates the MN extracellular voltage activity with spiking activity indicated with red triangles. The firing rates were converted into 25 ms window widths and smoothened with a gaussian filter (middle). Synthetic EMG activity was generated from recorded spiking activity by convolving gaussian wavelets (right). Grey shading highlights detected bursts of firing rate activity.

### Spiking Neural Network

The SNN model simulated ipsilateral rodent ankle flexor’s mono- and di-synaptic stretch and stretch velocity afferent reflexes. Proprioceptive afferents innervated the TA MN, GABA, Ia inhibitory, and V2a interneurons (88, 89). Ia inhibitory interneurons (IaINs) receiving Ia and II afferent inputs of the flexor and extensor muscles were reciprocally inhibited (79). GABA INs applied presynaptic inhibition to excitatory inputs of the TA MN (69, 90). V2a INs received flexor II afferent inputs and applied tonic excitation to TA MNs (77, 91). TA MNs received monosynaptic excitation from flexor Ia afferents (81, 92). Refer to Fig. 1 A for an illustration of the entire network and figures 9 and 10 for tonic bursting neural firing. IaINs were modelled as conductance-based LIF neurons receiving excitation from Ia and II afferent fibres and inhibition from opposing IaINs, see equation (4). *I*_*syn*_ is the cumulative synaptic current from excitatory and inhibitory components. IaIN parameters in table 2 were set to match experimental results (62, 68).

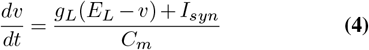

GABA presynaptic inhibitory INs and V2a INs were modelled as conductance-based adaptive exponential (AdEx) LIFs (84). GABA parameters (80, 83) and V2a parameters (75, 77, 78) were set as per experimental results. AdEx equation was defined per equation (5). GABA and V2a IN parameters were set per table 2. V2a IN g_*L*_ was reduced by 40% during SCI experiments and 15% during BWS experiments (85).

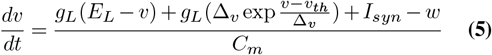

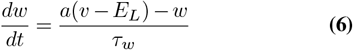

TA flexor MNs were modelled as exponential LIFs receiving excitation from TA Ia fibres, V2a INs, inhibition from GM originating IaINs and presynaptic inhibition from GABA INs (38, 62, 93). The equation for TA MNs was the same as equation (5) where *w* = 0. Parameters in table 2 were set to best estimate experimental results (68, 94, 95). To simulate serotonin agonist response, MN g_*L*_ was reduced by 40% during SCI experiments and 15% during BWS experiments (62, 85, 96). See Fig. 9 for MN responses with varying stimulation pulse widths.

**Table 2.** Ia inhibitory, GABA, V2a IN and MN parameters.

| Parameter | IaIN | GABA | V2a | MN |
| --- | --- | --- | --- | --- |
| N | 196 | 196 | 196 | 169 |
| $C_m$ | 31.13 pF | 100 pF | 45 pF | 162 pF |
| $E_L$ | -70 mV | -70 mV | -53 mV | -70 mV |
| $V_{th}$ | -50 mV | -50 mV | -42 mV | -50 mV |
| $V_{rest}$ | -65 mV | -62.3 mV | -47 mV | -65 mV |
| $g_L$ | 5 nS | 1.2 nS | 1.2 nS | 27 nS |
| $\Delta_v$ | — | 2 mV | 0.5 mV | 0.05 mV |
| $a$ | — | 2 nS | 2 nS | — |
| $\tau_w$ | — | 20 ms | 55 ms | — |

### Synapses

Alpha and exponential conductance synapses were used to describe inhibitory and excitatory synapses, respectively, see table 3 and Fig. 1 D and E. The reversal potential of excitatory synapses was set to 0 mV, while inhibitory synapses were set to the target neuron reverse potential. IIfibre synapse weights were scaled by a factor of 0.33 to simulate the effect of smaller axon size (62, 97). Synaptic connections, with the exclusion of GABA, were determined by probabilities specified in table 4. Axon synapses included a 2±0.3 ms delay accounting for diameter variability (98). GABA connections were predetermined by index rules dependent on the experiment condition; see Fig. 1 C. GABA connections were tuned to match previous baseline responses (59, 62). SCI condition GABA connections were increased by 160% as seen in flexor MNs after SCI transection (69).

**Table 3.** Alpha and exponential synapse and GABA spillover parameters. Alpha synapses were used for inhibitory connections while exponential synapses were used for excitatory connections.

| Parameter | Value |
| --- | --- |
| $\tau_{exc}$ | 0.25 ms |
| $\tau_{inh, rise}$ | 2 ms |
| $\tau_{inh, decay}$ | 4.5 ms |
| $\tau_p$ | 20 ms |

**Table 4.**
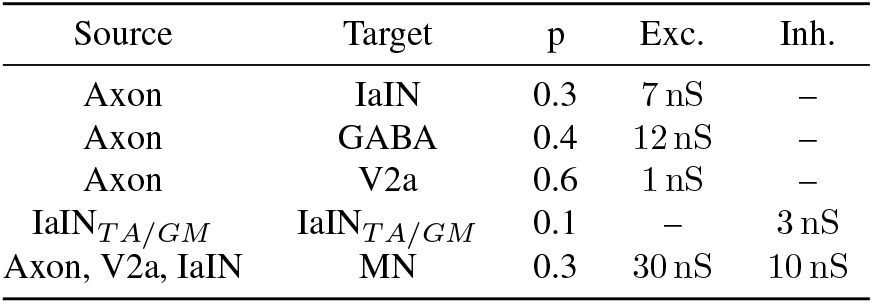
Synapse connection probabilities (p) and synaptic conductance by source and target neurons. Excitatory (exc.) and inhibitory (inh.) synaptic conductance apply to target neurons.

| Source | Target | $p$ | Exc. | Inh. |
| --- | --- | --- | --- | --- |
| Axon | IaIN | 0.3 | 7 nS | – |
| Axon | GABA | 0.4 | 12 nS | – |
| Axon | V2a | 0.6 | 1 nS | – |
| IaIN<sub>TA/GM</sub> | IaIN<sub>TA/GM</sub> | 0.1 | – | 3 nS |
| Axon, V2a, IaIN | MN | 0.3 | 30 nS | 10 nS |

Presynaptic inhibition was a multiplicative gain, scaling synaptic weights of each excitatory connection to the TA MN population (72, 80, 99, 100). GABA spillover was modelled as a linear decrease in release factor, p, see equation (7). *β* determined the strength of the inhibition and set to a value of 0.4. C_*GABA*_ was considered a unitless value for local GABA concentration (101).

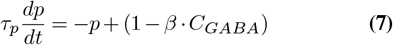

Poisson distributed neural groups simulated subthreshold stimulation with user-specified input rates. Each stimulation input was connected to three afferent fibres (Fig. 1C). To validate the SNN model, an EMG signal (Fig. 1F) was generated using convolutions of representative motor unit action potentials using the same parameters as experimental and previously validated computational models (59, 105).

## Results

A spinal reflex recruitment protocol validated the interactions between neural groups, synapses, and ES and afferent axons in a static environment. Early(ER), middle(MR), and lateresponse (LR) latencies were defined as 1 ms, 4 ms, and 7 ms respectively (106). The protocol was adapted from previous animal and simulation experiments and qualitatively assessed the SNN model validity (59, 102). The shape, latency, and modultation of evoked responses were qualitatively comparable to in-vivo measurements (Fig. 2A). Increased stimulation intensities led to the recruitment of efferent fibres, suppressing MR and LR amplitudes, matching previously reported simulations and experimental data.

**Fig. 2.**
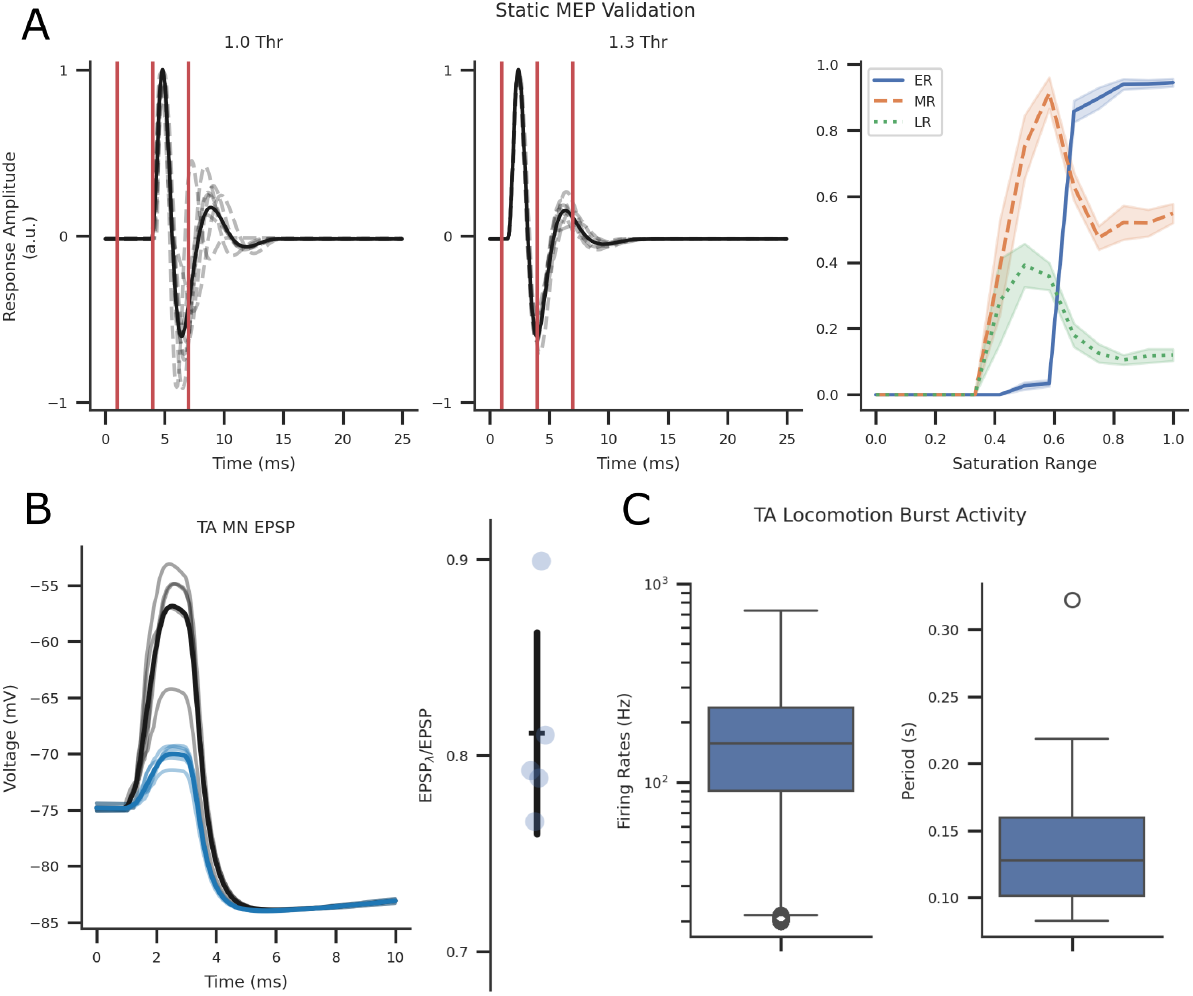
Static (n = 7) and dynamic (n = 9) qualitative and quantitative validation of the SNN model. **A** Motor evoked potentials (MEPs) at stimulation intensities ranging from 0 – 600 µA and 0.5 ms pulse width with 1 s between each pulse (59, 102). Results qualitatively match reported findings. **B** Modulation of excitatory post-synaptic potential in control (black trace) and GABA IN stimulation (blue trace) experiments. Fifteen 1 ms pulses at 50 Hz with a 45 ms delay stimulated GABA INs before delivering afferent fibre stimulation at 1.1x threshold (80). The ratio of maximum peaks was within the range of experimental results. **C** TA burst firing rates and periods during baseline locomotion were within the the range of experimental results. (103, 104).

Additionally, GABA interaction validation was performed by adopting electrophysiological evoked response experiments in intracellular recordings of the TA MN (Fig. 2B). To simulate the photostimulation of GABA neurons, a threshold current pulse was injected using the same pulse durations, frequencies, and delays from experimental protocols (80). The ratio of extracellular potential during control and photostimulation conditions was within range of experimental results. The SNN model was further verified within a dynamic setting (Fig. 2C) using previously validated afferent fibre firing rate profiles during locomotion (62). The MN firing rates were recorded and processed with the same window size (4 ms) as experimental methods in rodents and validated by comparing TA MN burst firing rates and periods during locomotion (103, 104). The burst activity during locomotion fell within the range reported in experimental data.

Statistical analysis on flexor activity after outlier removal (SCI, SCI_5-HT+ES_) during each phase presented a non-normal distribution in the swing and stance phases (p < 0.05). Swing phase flexor activity significantly differed between all simulated conditions with the exclusion of baseline – BWS_5-HT+ES_ and BWS_ES_ pairs. Stance phase flexor activity was significantly different in all simulated conditions except baseline – BWS_ES_ pairs. See Fig. 3 for TA MN activity distributions in both swing and stance phases. Box-and-whisker plots for GABA INs and V2a INs can be found in figures 11 and 12. Simulated TA MN expression during baseline step cycles was unaffected by stimulation amplitudes but minorly affected by frequencies, refer to Fig. 5. Though, GABA and V2a INs firing rates were scaled according to stimulation frequency. Note 20 Hz and 40 Hz at 10 mV stimulation intensities resulted in the same firing patterns.

**Fig. 3.**
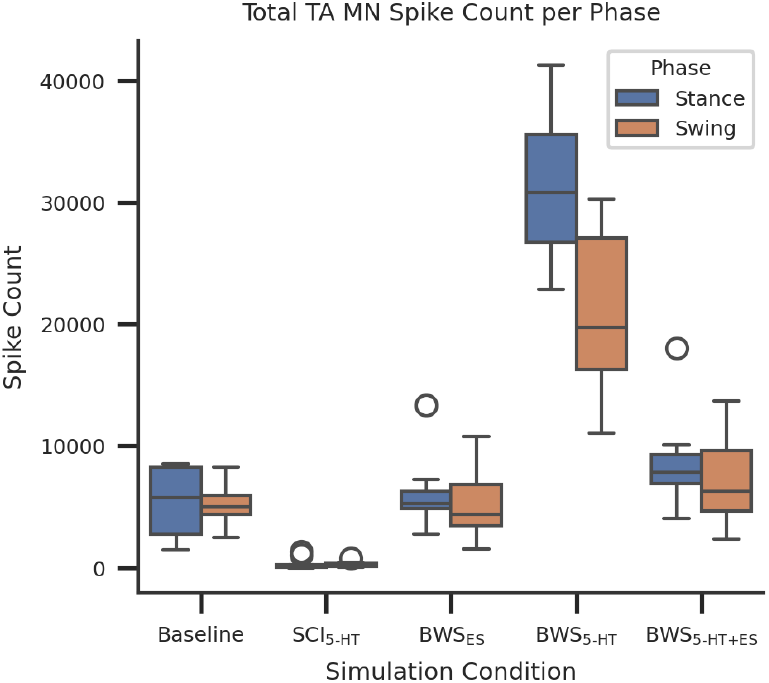
Box-and-whisker plots of 9-steps during baseline and simulated conditions. All swing flexor activity was non-normal and significantly different with the exception of Baseline to BWS_ES_ and BWS_5-HT+ES_. All stance flexor activity was significantly different with the exception of Baseline to BWS_ES_.

During baseline stepping, most variation occurred within the swing stages and transition between the swing and stance phases of the gait cycle, see Fig. 4. The TA MN population firing rates between baseline and SCI conditions were significantly different. Deterministically scaling the GABAergic connectivity to flexor MNs by 1.6 reduced mean firing rates by 53.6 Hz and increased the coefficient of variation by a factor of 9.2.

**Fig. 4.**
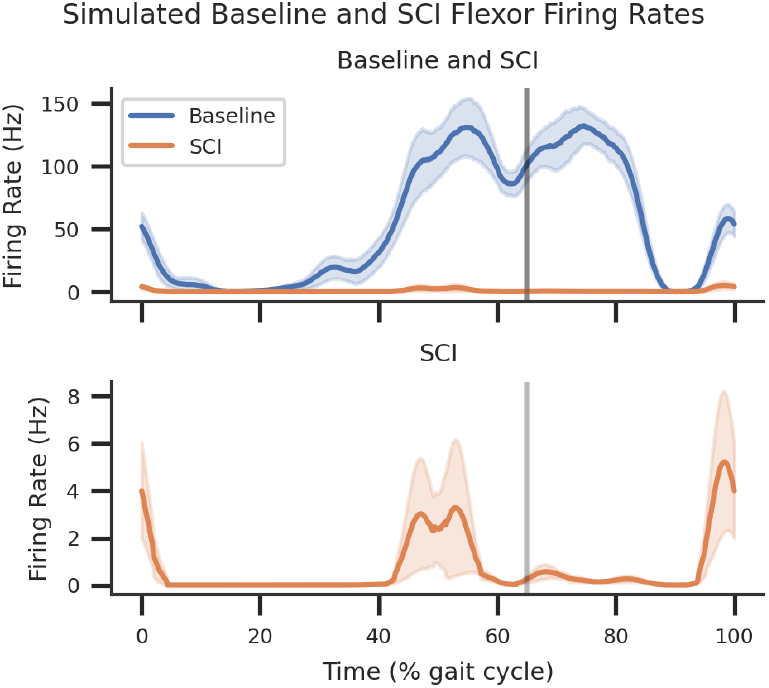
Average and standard deviation plots of 9-steps during baseline and simulated SCI conditions. The vertical grey line separates the stance (left of the grey line) and the swing (right of the grey line) phases, estimated at 65% of the gait cycle (67).

**Fig. 5.**
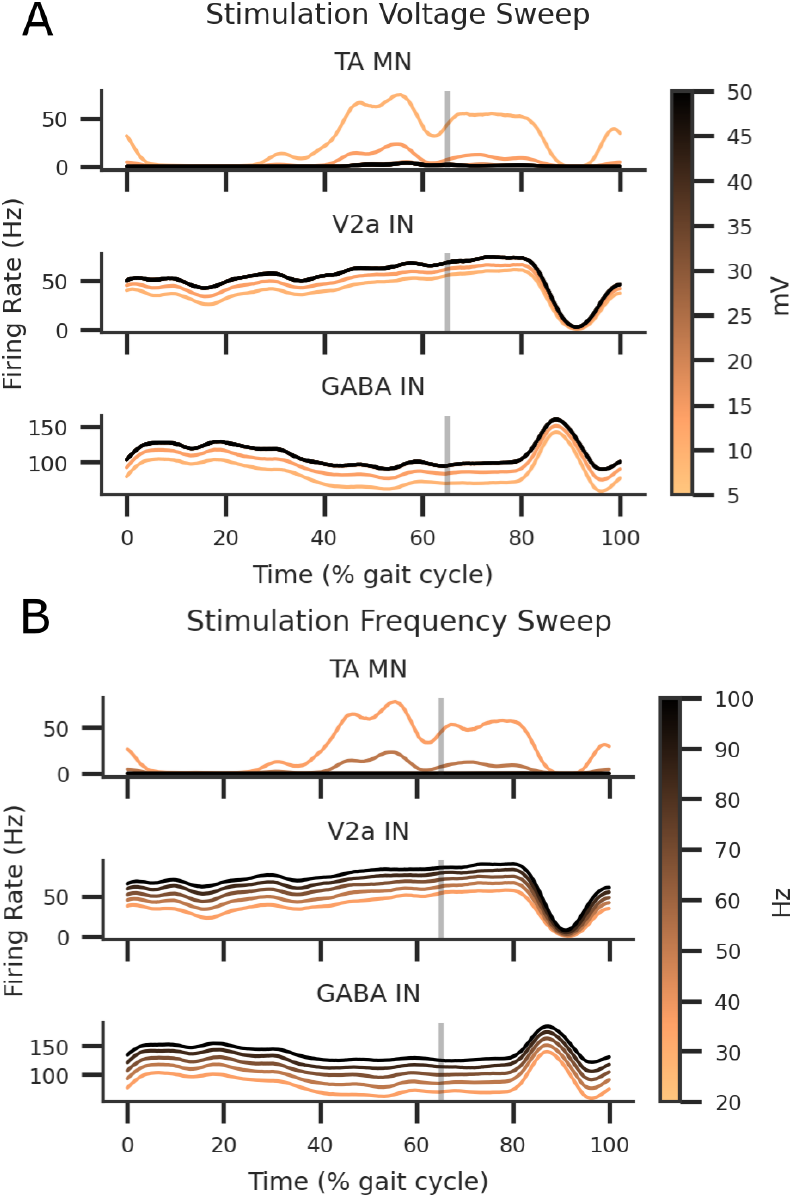
ES intensity and frequency sweeps using poisson inputs to flexor and extensor afferent axons. **A** Stimulation intensity sweep applied at 5, 10, 20, 30, 40, and 50 mV at a 40 H_z_ frequency. **B** Stimulation frequency sweep applied at 20, 40, 60, 80, and 100 H_z_ at 10 mV amplitude.

While the GABA IN firing rates were the same between simulated baseline and SCI settings, increased GABAergic connections resulted in more frequent presynaptic inhibition activity on TA MN populations (Fig. 3). GABA IN firing rates were increased while receiving ES inputs. V2a IN firing rates were equal between baseline and SCI since it did not receive GABA IN synapses. Simulating SCI serotonergic agonist activity by reducing the membrane conductance of V2a INs and MNs only slightly increased firing rates in MNs. This effect was abolished when combined with ES, see figures 3 and 6. Applying 5-HT+ES increased V2a IN and GABA IN activation, reducing the MN excitation to below SCI. In combination with the SCI condition, ES resulted in the same MN firing rate expression as 5-HT+ES.

**Fig. 6.**
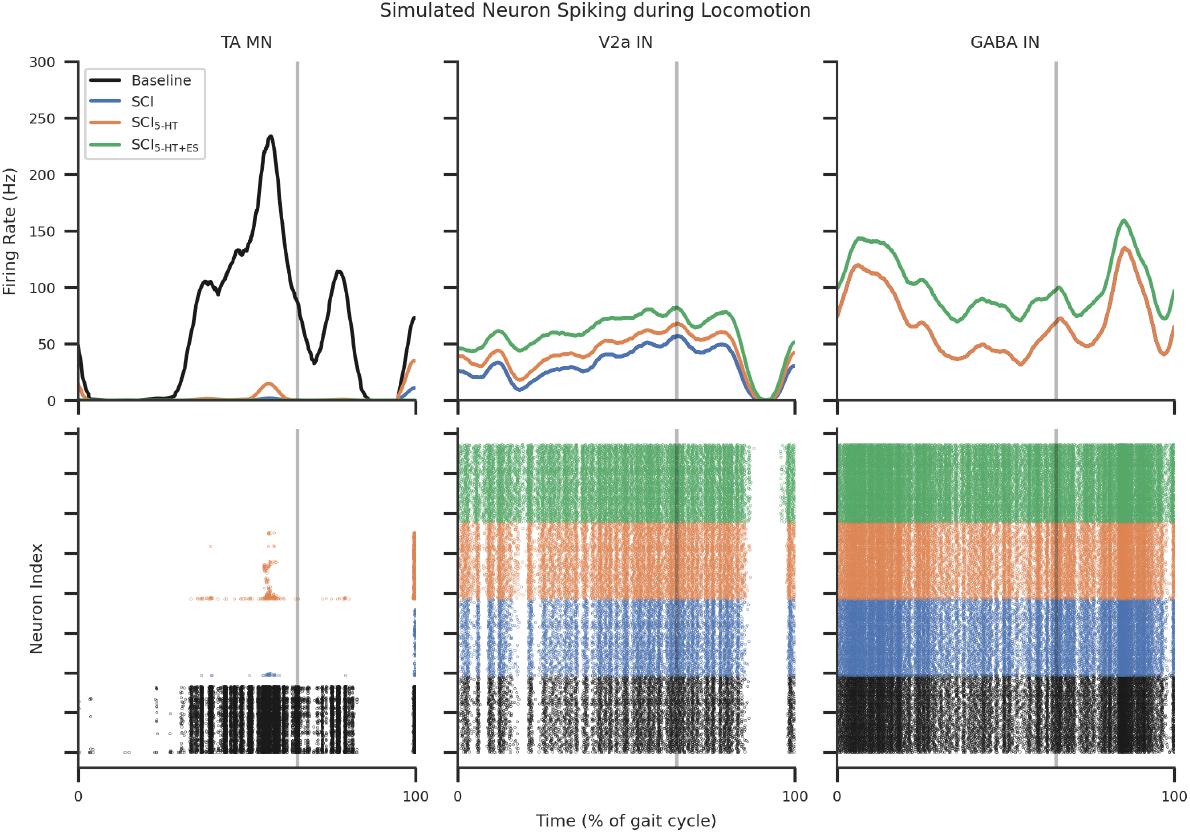
MN, V2a IN, and GABA IN spiking activity during an example step. The effect of SCI and SCI while receiving serotonergic agonists, (5-HT), and the combination of 5-HT and ES (5-HT+ES) were compared. The top row illustrates the population firing rates and the bottom row shows the individual neuron spiking activity during the gait cycle.

BWS locomotion with SCI increased overall flexor activity to averages greater than the baseline condition. This was further amplified with the introduction of 5-HT. Applying ES without 5-HT smoothened the output of MN activity, returning MN activations to baseline. Combining 5-HT and ES further increased peak activity during the swing phase and reduced activations during stance phases, see figures 3 and 7.

**Fig. 7.**
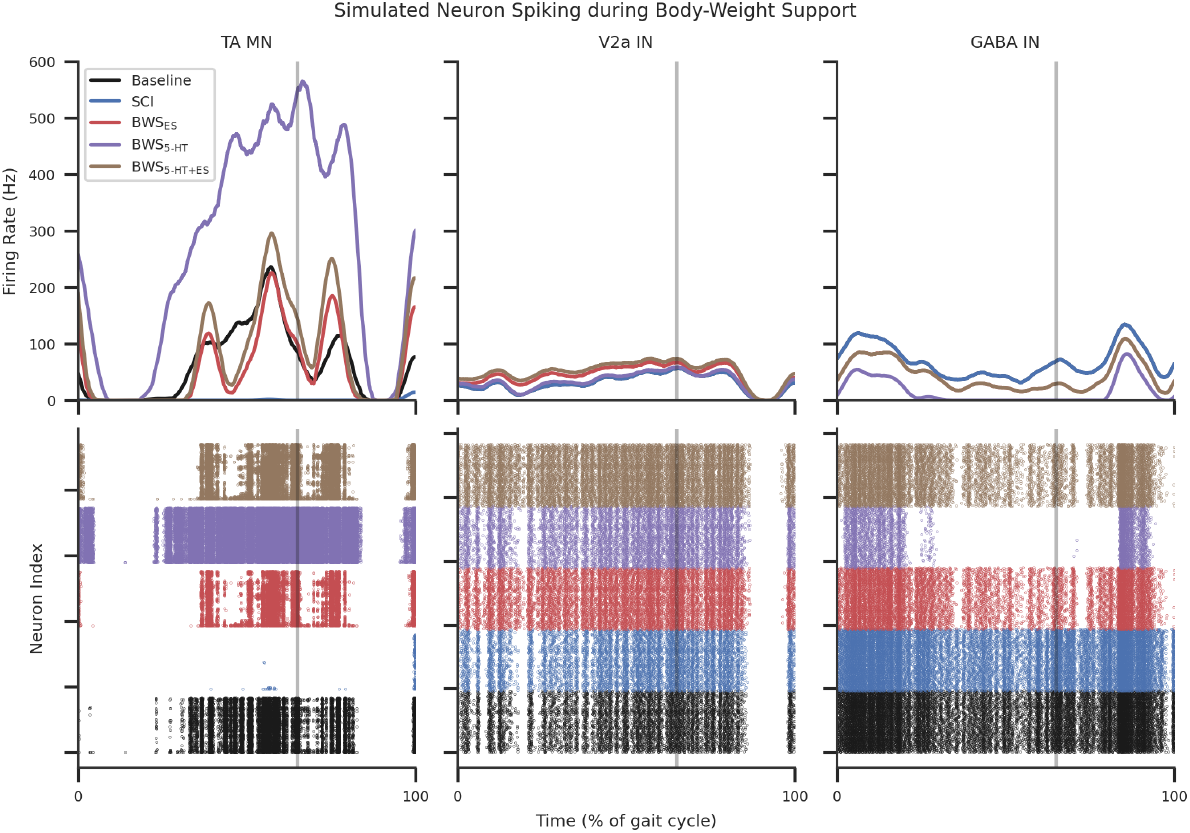
MN, V2a IN, and GABA IN spiking activity during an example step during simulated BWS locomotion. The effect of BWs while receiving serotonergic agonists (5-HT) and the combination of 5-HT and ES (5-HT+ES) were compared. The top row illustrates the population firing rates and the bottom row shows the individual neuron spiking activity during the gait cycle. Descreased firing rates from extensor afferents increased excitation in V2a INs and MNs by reducing the effect of inhibition.

The reduction in flexor afferents from BWS reduced the 5HT-modulated V2a IN activity towards SCI levels; these effects were reversed with ES modulation. GABA IN activity was reduced due to BWS simulated stance EMG reduction. Introducing ES returned GABA IN activity closer to the baseline. GABA activity was equivalent when comparing BWS_ES_ and BWS_5-HT+ES_ since no modulation was applied from serotonin, refer to figures 3 and 7.

## Discussion

A biologically constrained SNN model of the flexor reflex circuit was designed to investigate the integrative mechanisms between sensory and neuromodulation inputs to the spinal cord. Analysis of the stance and swing phases of 9 steps during simulated SCI activity revealed serotonergic agonists as a method to excite V2a INs and TA MNs after SCI. Applying unspecific ES to proprioceptive afferent axons amplified the effects of reciprocal inhibition, further accentuating excitatory peaks and inhibitory valleys. Moreover, simulating BWS locomotion decreased presynaptic inhibition, which reestablished TA MN firing rates. Applying ES during BWS locomotion resulted in a smoother MN activation profile, returning swing and stance firing rates to the baseline level.

Historically, ES has been applied for chronic pain management and spasticity reduction (65, 107, 108). The activation pathways of spinal cord stimulation for pain management are understood to be via large-diameter dorsal column and root fibres that carry propriosensory, mechanosensory, and nociceptive information (109). GABA INs activate and depress afferent nociceptive signals by antidromic activation of the dorsal column at frequencies, electrode positions, and stimulation amplitudes similar to that of ES for sensorimotor recovery (110, 111). Similarly, ES applied for spasticity reduces the excitatory inputs to MNs through the proprioceptive inhibitory pathways (112, 113). Yet, literature in SCI motor recovery places an intense focus on excitation (31, 114, 115).

Given the heightened inhibitory state of the injured spinal cord, it seems intuitive to return excitation to depressed neurons. However, results from this study suggest activating the spinal cord with the same proposed mechanisms as pain and spasticity modulation equally activate inhibitory pathways, strengthening an already maladapted inhibition dominant circuit (69, 116). A more refined and nuanced approach needs to be considered in order to return the required balance of excitation and inhibition to allow phasic activity to propagate. Results in this study suggest that appropriate sensory information must be provided to drive flexor network plastic adaptation towards a less inhibited and more task-specific tuned state. Tonically depressing or exciting, the spinal circuits do not provide the necessary sensory information to provide that plastic tuning (117, 118). This explains the requirement of propriosensory information for locomotor recovery after SCI (32–34).

The normal sensory processing occurring within the injured spinal cord becomes more stochastic and lacks the necessary bias required to perform the task (106). As a result, pre-motor polysynaptic connections play a more active role in the expression of muscle tone and activity (119). By establishing an appropriate balance in excitation and inhibition, repetitive sensory information can reinforce appropriate synaptic adaptations towards a more functional spinal state (60, 120). Results in this study show, for the first time, the synaptic mechanisms at which this can be accomplished and provide an understanding of why BWS treadmill training works (121–123).

The greater deviation in normalised firing rate indicates greater variation in neuronal populations after SCI (figures 4 and 8). This may be due to the lack of necessary excitation required to inhibit and excite in phase with the incoming sensory signals (106). Thus, increasing the excitability of MNs and premotor excitatory neurons and reducing the effect of stance phase inhibition by BWS can assist with reducing inhibitory effects on flexors (124–127). After reaching a suitably excitable state where phasic step information can propagate in a timely manner, sub-threshold ES could provide the necessary smoothening process to synergistically reinforce relevant pathways without too much inhibition or excitation (62, 128, 129).

**Fig. 8.**
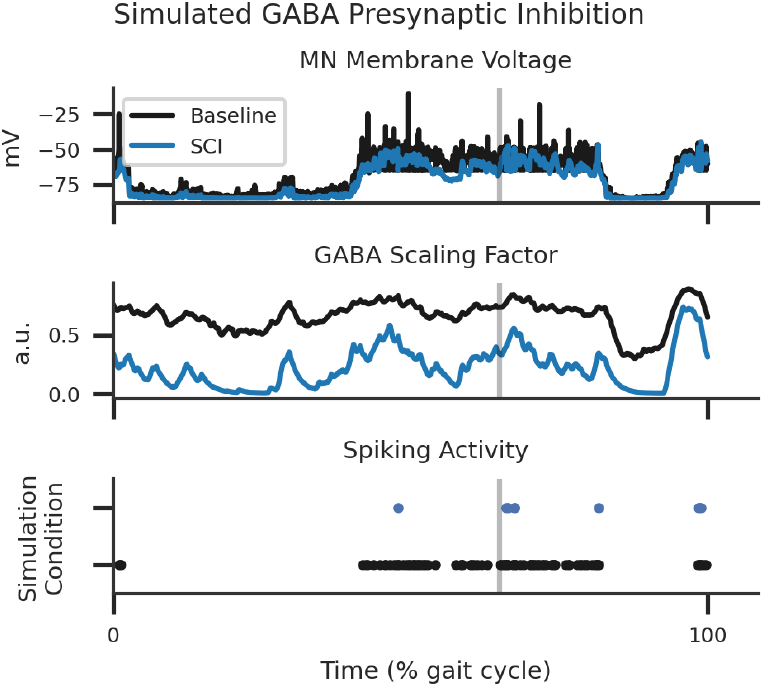
SCI simulation of a single motoneuron receiving presynaptic inhibition by local concentration of GABA transmitters during a single step. Note the reduced number of spike events due to GABA inhibition.

Computational studies such as this are limited in their ability to generalise due to the estimates and tuning that are required to generate the model itself. The simulated flexor reflex loop’s SNN architecture is simplistic compared to the complex bidirectional information exchange between the contralateral sides (130, 131). Though the cells were modelled from experimental data, there are errors and missing variables within experiments that have a carry-on effect on computational models. This study utilised LIF and AdEx equations to reduce computational burden and improve simulation runtime speeds. Though previous efforts have incorporated the same approach (59, 62), mathematical approximations of firing patterns limit the generalisability (132, 133). However, even with simple estimations, a computational model could provide new hypotheses to the inner workings of neurological systems and unlock novel recovery protocols (111, 134).

Future studies could include the experimental verification of these findings via electrophysiological or genetic ablation experiments in rodent models in BWS neuromodulated locomotion contexts. Moreover, extending the model enable investigation of previous electrophysiological results that uncovered correlations between the appearance of long-latency polysynaptic potentials and recovery of locomotion in spinal rats (35, 57, 102, 128, 135, 136). Re-emergence of uninterrupted late-response polysynaptic potentials may be the expression of increased excitability in local spinal networks, re-balancing the inhibitory dominant pre-motor circuity (106, 119). Functional recovery may be mediated by increased magnitude in polysynaptic activity, compensating for the loss in direct excitation (60). Finally, extending the computational model to include neuroplastic dynamics could uncover the relationship between neuromodulation and neuroplastic adaptations (101, 137, 138). Investigating these effects within an extended biologically constrained computational model would be worthwhile.

## Conclusions

The development of a biologically constrained SNN has provided insights into the mechanistic basis of sensory and neuromodulatory integration after SCI. Simulations suggest that BWS locomotion in conjunction with ES returns phasic flexor coordination in an inhibition-dominant environment. Without BWS, serotonergic agonists increased excitation to enable sensory-driven flexor rhythmic activation. The spinal cord requires a balance of excitation and inhibition to enable the correct phasic modulation of sensorimotor pathways. The work described in this chapter provides a potential explanation for why BWS locomotor training works with neuromodulation.

## ACKNOWLEDGEMENTS

We would like to thank V. Reggie Edgerton and Parag Gad (University of California, Los Angeles) for the invaluable scientific discussions which led to the conception of the experiment. We would also like to thank Bryce Vissel (St. Vincent’s Hospital and University of New South Wales Sydney) for having initiated SCI research at University of Technology Sydney (UTS). Thank you to Howe Zhu (UTS, University of Sydney) for the feedback and support over the years. Finally, thank you to Matthew Gaston of the UTS eResearch team for providing support and system administration of the Interactive High Performance Computing (iHPC) facility. The computing power and stability of the iHPC system was paramount to running the experiment. Funding was provided by the Research Training Program (RTP) scheme from the Department of Education and Training, Australia.

