## Supplemental Table and Figures for "Biologically-Constrained Spiking Neural Network for Neuromodulation in Locomotor Recovery after Spinal Cord Injury"

### Supplementary Note 1: Supplementary Information

**Table 5.** TA and GM afferent axon tuning performance measured by Pearson correlation coefficient (CC) and mean absolute error (MAE). All correlations were significant ($p < 0.05$).

| Afferent | CC | MAE (Hz) |
|---|---|---|
| TA Ia | 0.99 | 3.07 |
| TA II | 0.99 | 2.38 |
| GM Ia | 1.00 | 3.13 |
| GM II | 0.99 | 2.12 |

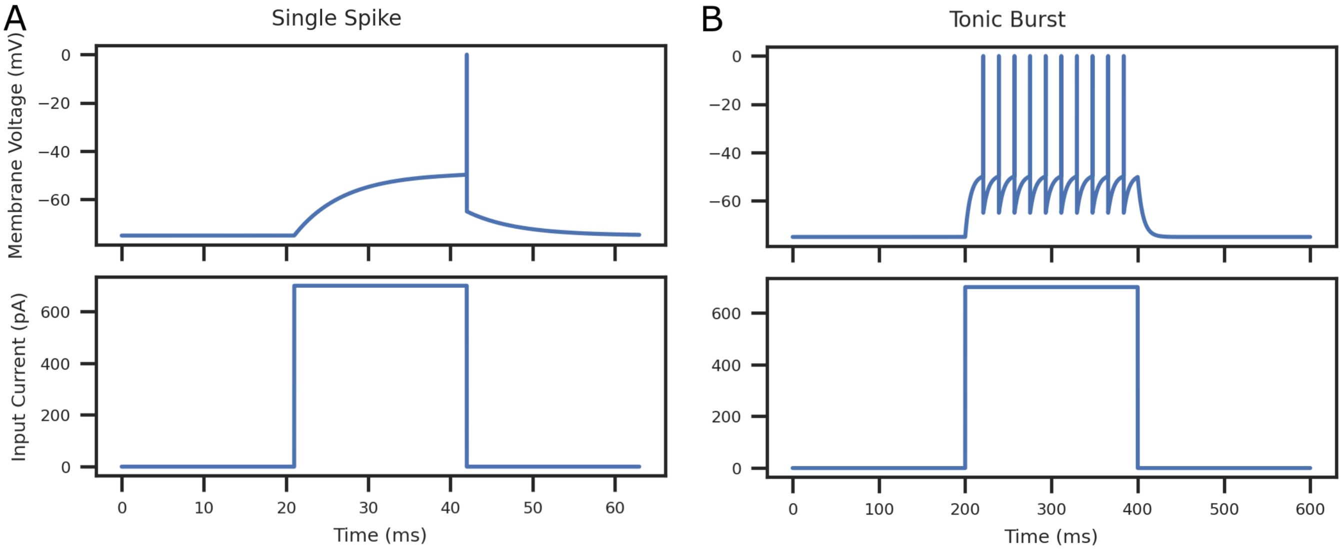


**Fig. 9.** Simulated AdEx LIF TA MN single spike (**A**) and tonic burst (**B**) response after receiving a $20\,\mathrm{ms}$ and $200\,\mathrm{ms}$ stimulation pulse at $670\,\mathrm{pA}$, respectively.

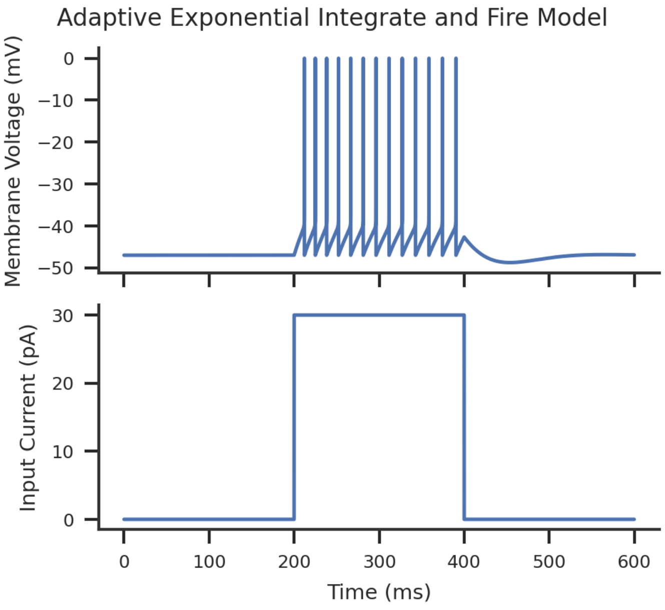


**Fig. 10.** Simulated AdEx LIF V2a IN tonic spiking response after receiving a $200\,\mathrm{ms}$ stimulation pulse at $30\,\mathrm{pA}$.

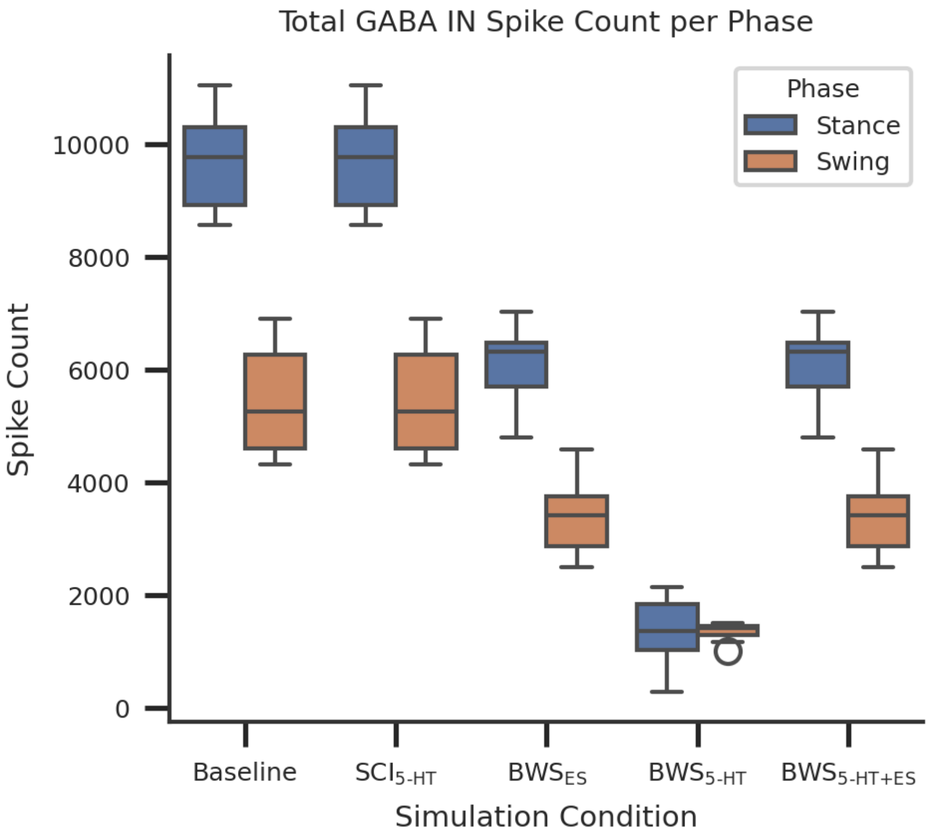


**Fig. 11.** Box-and-whisker plots of GABA IN spiking activity for 9-steps during baseline and simulated conditions. Shapiro-Wilk test returned a normal distribution for swing phase only. All stance flexor activity was significantly different after Wilcoxon signed-rank test with the exclusion of Baseline – $SCI_{5\text{-}HT}$ and $BWS_{ES}$ – $BWS_{5\text{-}HT+ES}$. All swing flexor activity was significantly different after Tukey HSD test with the exclusion of Baseline – $SCI_{5\text{-}HT}$ and $BWS_{ES}$ – $BWS_{5\text{-}HT+ES}$.

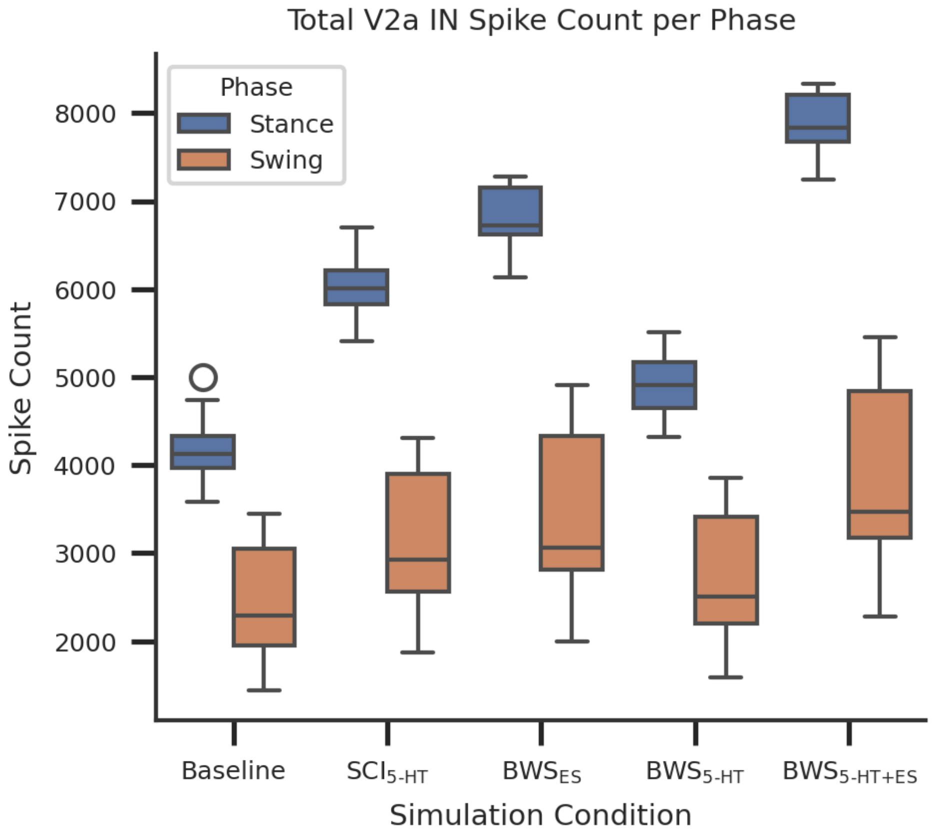


**Fig. 12.** Box-and-whisker plots of V2a IN spiking activity for 9-steps during baseline and simulated conditions. Shapiro-Wilk test returned a normal distribution for both phases. All stance flexor activity was significantly different after Tukey HSD test with the exclusion of $SCI_{5\text{-HT}}$ – $BWS_{5\text{-HT}}$. Baseline swing flexor activity was significantly different compared with $SCI_{5\text{-HT}}$, $BWS_{5\text{-HT}}$, and $BWS_{ES}$. Additionally, $SCI_{5\text{-HT}}$ swing flexor activity was significantly different compared with $BWS_{5\text{-HT}}$, $BWS_{ES}$, $BWS_{5\text{-HT+ES}}$. Finally $BWS_{5\text{-HT}}$ in the same activity was significantly different compared to $BWS_{5\text{-HT+ES}}$ after Tukey HSD test.
